# Mesoscale Modeling of a Nucleosome-Binding Antibody (PL2-6): Mono- vs. Bivalent Chromatin Complexes

**DOI:** 10.1101/607218

**Authors:** Christopher G. Myers, Donald E. Olins, Ada L. Olins, Tamar Schlick

**Affiliations:** Department of Chemistry, New York University, 100 Washington Square E, New York, NY, 10003, United States; Department of Pharmaceutical Sciences, College of Pharmacy, University of New England, 716 Stevens Avenue, Portland, ME 04103, USA; Courant Institute of Mathematical Sciences, 251 Mercer St, New York University, New York, NY, 10012, United States; New York University-East China Normal University Center for Computational Chemistry at New York University Shanghai, Room 340, Geography Building, 3663 North Zhongshan Road, Shanghai, China

## Abstract

Visualizing chromatin adjacent to the nuclear envelope (denoted “epichromatin”) by *in vitro* immunostaining with a bivalent nucleosome-binding antibody (termed monoclonal antibody PL2-6) has suggested a distinct and conserved chromatin structure. Moreover, different staining patterns for chromatin complexed with the monovalent “Fab” fragment of PL2-6, compared to the bivalent form, point to distinct binding interactions. To help interpret antibody/chromatin interactions and these differential binding modes, we incorporate coarse-grained PL2-6 antibody modeling into our mesoscale chromatin model and analyze interactions and fiber structures for the antibody/chromatin complexes in open and condensed chromatin, with and without linker histone H1 (LH). Despite minimal and transient interactions at physiological salt, we capture differential binding for monomer and dimer antibody forms to open fibers, with much more intense interactions in the bivalent antibody/chromatin complex. For these open “zigzag” fiber morphologies, differences result from antibody competition for peptide tail contacts with internal chromatin fiber components (nucleosome core and linker DNA). Antibody competition results in dramatic conformational and energetic differences among *monovalent*, *bivalent*, and *free* chromatin systems in the parental linker DNA / tail interactions. These differences in binding modes and changes in internal fiber structure, driven by conformational entropy gains, help interpret the differential staining patterns for the monovalent versus bivalent antibody/chromatin complexes. More generally, such dynamic interactions which depend on the complex internal structure and self-interactions of the chromatin fiber have broader implications to other systems that bind to chromatin, such as linker histones and remodeling proteins.

**STATEMENT OF SIGNIFICANCE:** Using mesoscale modeling, we help interpret differential binding modes for antibody/chromatin interactions to elucidate the structural details of “epichromatin” (chromatin adjacent to the nuclear envelope), which had been visualized to produce different staining patterns for monovalent and bivalent forms of the PL2-6 antibody. To our knowledge, this is the first application of such a coarse-grained computational antibody model to probe chromatin structure and mechanisms of antibody/chromatin binding. Our work emphasizes how antibody units compete with native internal chromatin fiber units (histone tails, nucleosome core, and linker DNA) for fiber-stabilizing interactions and thereby drive differential antibody binding for open zigzag chromatin fibers. Such competition, which dynamically alters internal chromatin structure upon binding, could be relevant to other chromatin binding mechanisms such as those involving linker histones or chromatin remodeling proteins.

## INTRODUCTION

Epichromatin was hypothesized to be a distinct conformation of chromatin when a unique staining pattern was captured with antibody molecules beneath the interphase nuclear envelope (NE) of human cells U2OS and HL-60/S4 (**?**). Specifically, the *bivalent* mouse antibody PL2-6 produced a ring-like staining pattern on formaldehyde-fixed and detergent permeabilized cells (Fig. 1A). In contrast, the *monovalent* Fab form of PL2-6 stains chromatin throughout the fixed and permeabilized cell nuclei (Fig. 1A). These staining patterns suggest different binding modes between each form of the antibody and chromatin. Demonstrating the conservation of these staining patterns across diverse eukaryotic animal and plant chromatin, Olins and Olins (**?**) proposed the “epichromatin hypothesis”. Namely, they suggested that epichromatin may reflect a unique evolutionary conserved conformation of chromatin that facilitates an interaction with the nuclear envelope (NE) (**?**).

**Figure 1:**
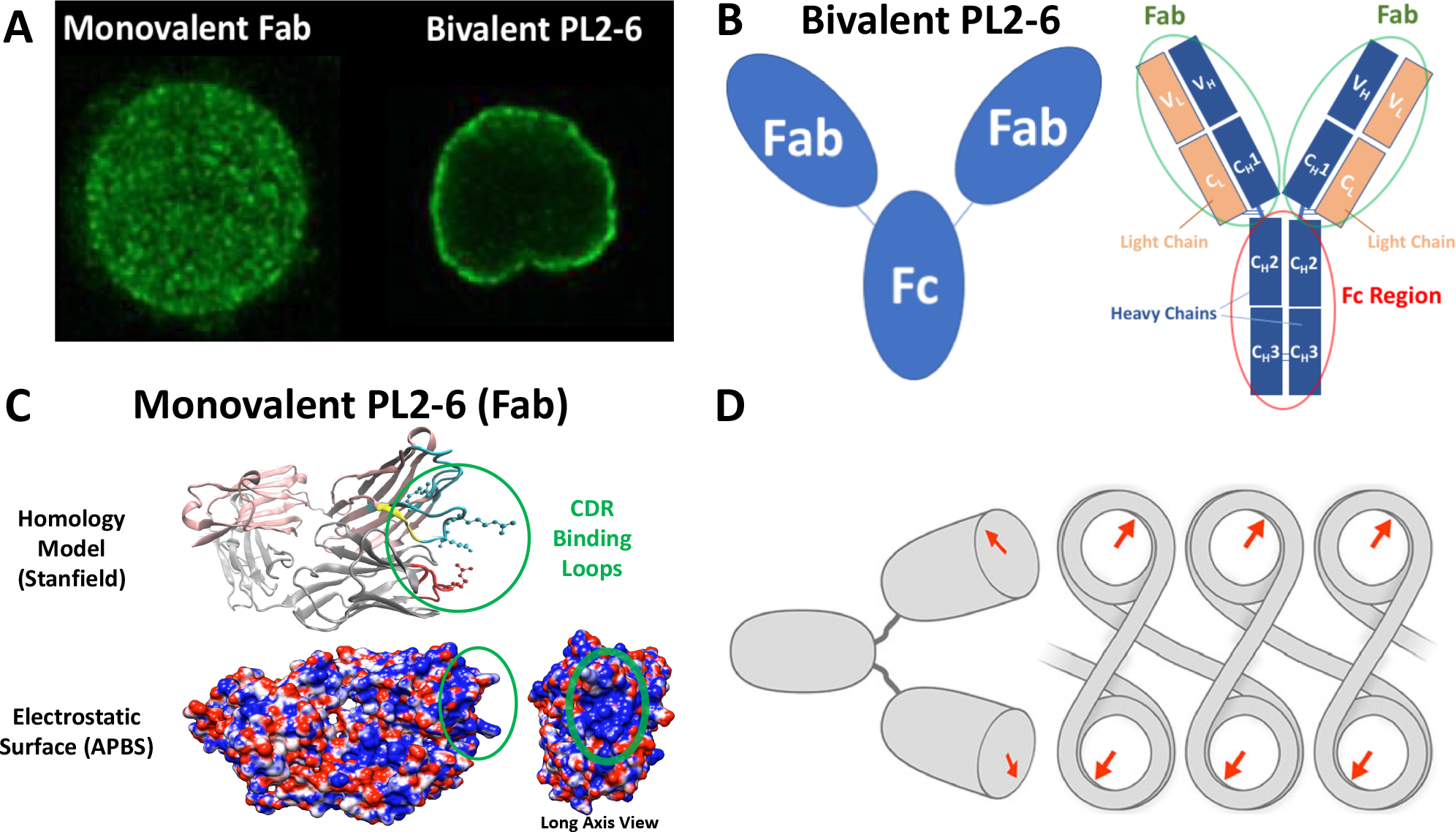
Structures and experiments demonstrating differences between monovalent and bivalent PL2-6 antibody binding to chromatin. (A) Differential staining patterns observed between two forms of PL2-6 which suggest differential binding modes to chromatin for each form. Monovalent PL2-6 ‘Fab’ stains across entire cell nuclei, whereas the bivalent PL2-6 produces a ring-like staining pattern localized near the nuclear envelope. (**?**). (B) Left: Cartoon depiction of IgG structure with two ‘Fab’ regions (circled in green) connected by Fc connector region (circled in red). Right: Schematic representation of heavy (dark blue) and light chains (orange) comprising the Fab and Fc regions, consisting of conserved (*C_H_* 1, *C_H_* 2, *C_H_* 3, *C_L_*) and variable (*V_H_*, *V_L_*) regions. Adapted from *Antibody Molecular Structure* (**?**). (C) Fab subunit homology model provided by Robyn Stanfield used to calculate the ‘Fab’ electrostatic surface using Adaptive Poisson-Boltzmann Solver (APBS) software (**? ?**) at pH 7.0. Binding regions with positive charge distributions from arginine residues are circled in green. (D) Potential mechanism for bivalent PL2-6 binding to dinucleosomes of chromatin at acidic patches by alignment along the dyad axis described by Olins and Olins (**?**).

PL2-6 is a mouse monoclonal (mAb) antibody belonging to the subclass G of immunoglobin antibodies. Immunoglobin G (IgG) antibodies represent the most physiologically abundant subclass. The IgG structure, sketched in Figure 1B, consists of a “crystallizable fragment” connector region (Fc) joining together two Fab (“antigen binding fragment”) arms. Each Fab arm contains identical highly-conserved CDR (“complementarity determining region”) loops. These disordered loops facilitate the highly-specific binding to an antibody’s preferred epitope target (**?**). Due to the binding affinity of both Fab arms of the IgG, we refer to the full IgG structure as the “bivalent” form of PL2-6. A “monovalent” Fab form of PL2-6, consisting only of a single Fab molecule, is derived via papain digestion through cleavage of the Fab arms from the Fc connector. Both the bivalent PL2-6 (**?**) and (papain-derived) monovalent Fab have been shown to bind all mononucleosomes in solution (**?**).

Despite the absence of X-ray crystal structures for PL2-6, the sequence homology of the PL2-6 CDR loop regions with centromere protein C1 (CENP-C) (**?**) and latency-associated nuclear antigen (LANA) (**? ? ?**) suggests that PL2-6 binds the nucleosome within the “acidic patch”. The acidic patch is a negatively-charged region found on both sides of symmetric nucleosome core particle and defined by six amino acid residues of the H2A/H2B histones (**? ?**). The monovalent Fab antibody and the Fab arms of the bivalent PL2-6 are hypothesized to bind specifically to the acidic patch (**?**). We have obtained a sequence-based homology model (Robyn Stanfield, personal communication) (**?**) for the monovalent Fab structure of PL2-6 (Fig. 1B). We use this predicted structure to calculate a Fab electrostatic surface which is specific to PL2-6. The high density of positive charge found at the antibody binding end due to arginines in the CDR loops further supports the hypothesis that the Fa binds at the negatively-charged acidic patch. The acidic patch also regulates high-order chromatin structure via the histone H4 N-terminal tail domains’ interactions with neighboring nucleosome core particles (NCPs). Subsequently, this region is crucial for chromatin remodeling and has important implications for gene regulation and cancer biology (**?**).

**Figure 2:**
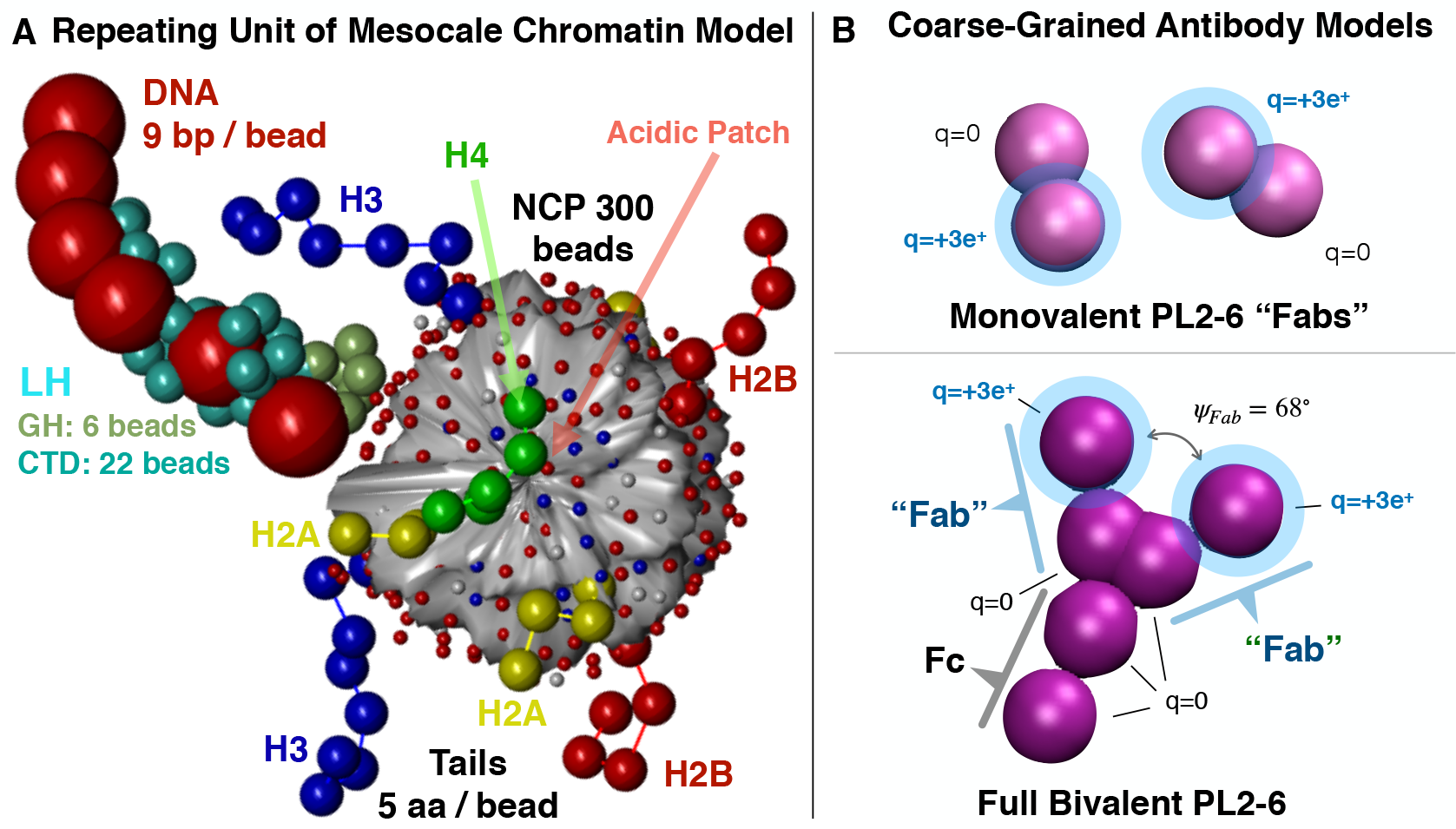
Mesoscale chromatin and coarse-grained antibody models. (A) Our chromatin mesoscale model repeating unit consists of a single nucleosome core particle (NCP) with 300 charged beads and flexible histone tails (5 amino acids per bead), joined to nearest neighbor NCPs by linker DNA (9 basepairs per bead). Linker histone (LH) consists of a C-terminal domain (CTD) and a globular head (GH). See details in Bascom *et al* (**?**). (B) Our coarse-grained rigid-body model for monovalent PL2-6 ‘Fab’ contains 2 beads. The bivalent PL2-6 model has two ‘Fab’ arms connected by a two-bead Fc connector region (with the same dimensions as a Fab), for a total of 6 beads. Positively charged spheres (highlighted in blue) are assigned Debye-Hückel-screened surface charges *q*_*F ab*_ = +3*e* at the binding end of the Fabs in both models.

The observed differences in staining patterns between the monovalent and bivalent forms of the PL2-6 antibody suggest distinct modes of chromatin binding for each form. Since each form shares the same binding region, geometric or internal variations in the chromatin structures might explain the preferential binding of the bivalent antibody. It has been hypothesized that the bivalent binding may occur via a cooperative two-step mechanism which results in a larger overall effective “association constant” (i.e., an “avidity”) for a particular chromatin geometry. As shown in Figure 1D, this proposed mechanism should depend on the symmetry and geometry of the bivalent IgG along its dyad axis, as well as on the chromatin conformation (**?**). The CDR binding loops of monovalent and bivalent antibodies likely interact with the acidic patch upon nucleosome binding and stabilize the resulting conformations. However, the internal interactions of the chromatin fiber could preemptively change the energetic landscape of chromatin binding, leading to differential binding. Moreover, monovalent and bivalent antibodies may follow different mechanistic pathways toward their final binding. Here we combine coarse-grained modeling of the PL2-6 antibody with our mesoscale chromatin model to elucidate differences in the structures and binding of chromatin complexes with bivalent versus monovalent PL2-6 antibodies. We show that differential binding of bivalent versus monovalent antibodies is driven by competition with native internal chromatin elements for tail interactions.

## MATERIALS AND METHODS

### Mesoscale Chromatin Model

We simulate chromatin fibers with our mesoscale chromatin model (Fig. 2A) (**? ? ?**) consisting of four components: coarse-grained beads of linker DNA, electrostatically-charged nucleosome core particles (NCPs), coarse-grained beads for flexible histone tails, and beads for H1 ‘linker histone’ (LH). Each NCP consist of 300 partial point charges which approximate the core electrostatic surface determined by applying our Discrete Surface Charge Optimization (DiSCO) algorithm (**? ?**) to the NCP crystal structure (**?**). NCPs are joined by linker DNA modeled as a modified worm-like chain of beads that each represent about nine basepairs. LH (based on rat H1.4 (**? ?**)) is modeled as a rigid body of 28 beads using the united atom model (**?**), with 22 beads for the C-terminal domain (CTD) and 6 beads for the globular head (GH) inserted on the dyad axis. For more details of our LH model see Luque *et al.* (**?**). In this study, we simulate fiber systems of 24 nucleosomes with and without LH. Equilibrium Monte-Carlo simulations with LH contain 24 linker histones at a total density *ρ* of one LH per NCP. All nonbonded interactions in the system are modeled with excluded-volume terms via 12-6 Lennard-Jones Van der Waals (VdW) potential and screened electrostatic Debye-Hückel energy terms

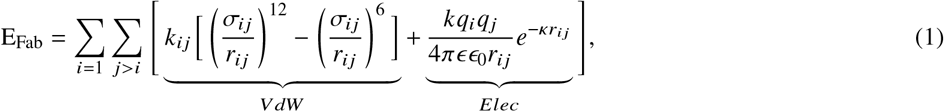

where σ_*ij*_ is the effective Lennard Jones van der Waals diameter of the two interacting beads in *nm* and *k*_*ij*_ is an energy parameter that controls the steepness of the excluded volume potential, *ϵ*_*r*_ = 88.9 is the relative dielectric of water, *ϵ*_0_ the permittivity of free space, and using a Debye screening length 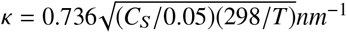 at *T* = 298*K* and monovalent salt concentration *C*_*S*_. There are 5 types of Monte Carlo (MC) moves in the chromatin system: a global pivot move, a configurationally-biased tail Rosenbluth regrow move, local rotation and translation moves for nucleosomes and linker DNA, and, for simulations including LH, translation of the LH CTD. All moves are accepted or rejected according to the Metropolis algorithm acceptance criterion with Boltzmann-weighted probabilities from the total system energy and temperature. For full details of the model see Bascom *et al* (**?**).

### Coarse-grained Antibody Model

We have devised and employed coarse-grained models (Fig. 2B) for both monovalent and bivalent PL2-6 antibody structures to simulate in conjunction with our mesoscale chromatin model. Such a coarse-grained bivalent antibody model is similar to previous models for IgG antibodies (**? ? ?**). To our knowledge, this is the first application of such a model to investigate antibody binding to chromatin or chromatin structure.

A homology model provided by Robyn Stanfield (personal communication) is used to perform electrostatic calculations using the Adaptive Poisson-Boltzmann Solver (APBS) software (**? ?**) and determine an electrostatic surface for the PL2-6 Fab (Figure 1C). Arginines within the CDR create a positive charge distribution at the binding end of the Fab subunit (green circles). This suggests that the energetics of Fab binding and stabilization with chromatin depend heavily on the electrostatic interaction between this positive charge distribution with the chromatin fiber via the negatively-charged acidic patch and the negatively-charged linker DNA. A coarse-grained electrostatic model of the antibody protein should thus capture significant aspects of fiber interactions leading to binding.

In both our coarse-grained antibody models (Fig. 2B), the Fab and Fc regions are modeled separately as two rigidly-connected spherical beads with volume exclusion and surface charges which interact with the beads of the chromatin fiber and the other antibodies, according to the energy terms

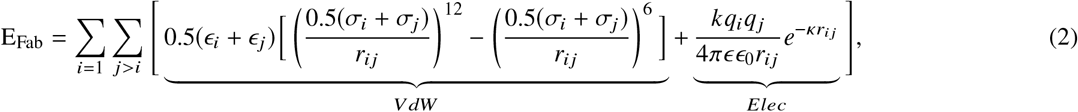

where *ϵ*_*i*_ = *k*_*b*_*T* and σ_*i*_ = 4.0 *nm* is the Lennard Jones van der Waals diameter for beads approximating the dimensions of the Fab and Fc regions, *q*_*i*_ is the charge of each antibody bead interacting with each other bead in the chromatin/antibody system according to their values of *ϵ*_*j*_, σ_*j*_, *q*_*j*_, and with electrostatic constants defined in Equation 1. The monovalent antibody consists of two rigidly-connected beads. The bivalent consists of a total of six beads of the rigidly connected Fab, Fab, and Fc regions. Each bead is assigned a surface charge *q*_*i*_ (monovalent: i=1,2; bivalent: i=1,2,…,6) which interacts with all other pairwise nonbonded charged beads of *q*_*j*_ and σ_*j*_ via Equation 2 and according to our previously-developed chromatin model. We employ a charge of *q*_*F ab*_ = +3*e* to approximate the charge of the Fab binding region containing the specific CDR loops and the three arginines mentioned above and thus assign *q*_*i*_ = *q*_*F ab*_ = +3*e* to a single bead in the monovalent Fab and to the end of the two-bead Fab groups in the bivalent PL2-6 (Fig. 2B, charged beads highlighted in blue). Thus, the entire bivalent antibody has a combined net charge of *q* = +6*e*.

Previous work has suggested that, although the IgG hinge region is flexible, the average experimental binding of IgGs can be captured using a rigid model with a fixed equilibrium “Fab” angle *Ψ* between the two subunits (**?**) (see Figure 5). Therefore, we model both the monovalent and bivalent antibodies as rigid molecules for computational simplicity at an equilibrium angle of Ψ_0_ = 68° used in previous IgG models (**? ? ?**). The bivalent model is constrained to be planar and rigid in the Fab and Fc connector regions which rotate freely in 3D space with orientations defined by three internal Euler angles α_*i*_, β_*i*_, γ_*i*_ (i = 1,2,…,.N for N antibodies). The antibodies move within the chromatin system according to two additional MC antibody translation and rotation moves, accepted or rejected according to the same Metropolis algorithm acceptance criterion with the Boltzmann-weighted probability from the total system energy and temperature. Thus, in bivalent and monovalent antibody/chromatin systems with LH, we employ a total of 7 Monte Carlo (MC) moves: the 5 chromatin moves mentioned above and these two additional antibody moves. In the absence of LH, there is one less relevant move.

**Figure 3:**
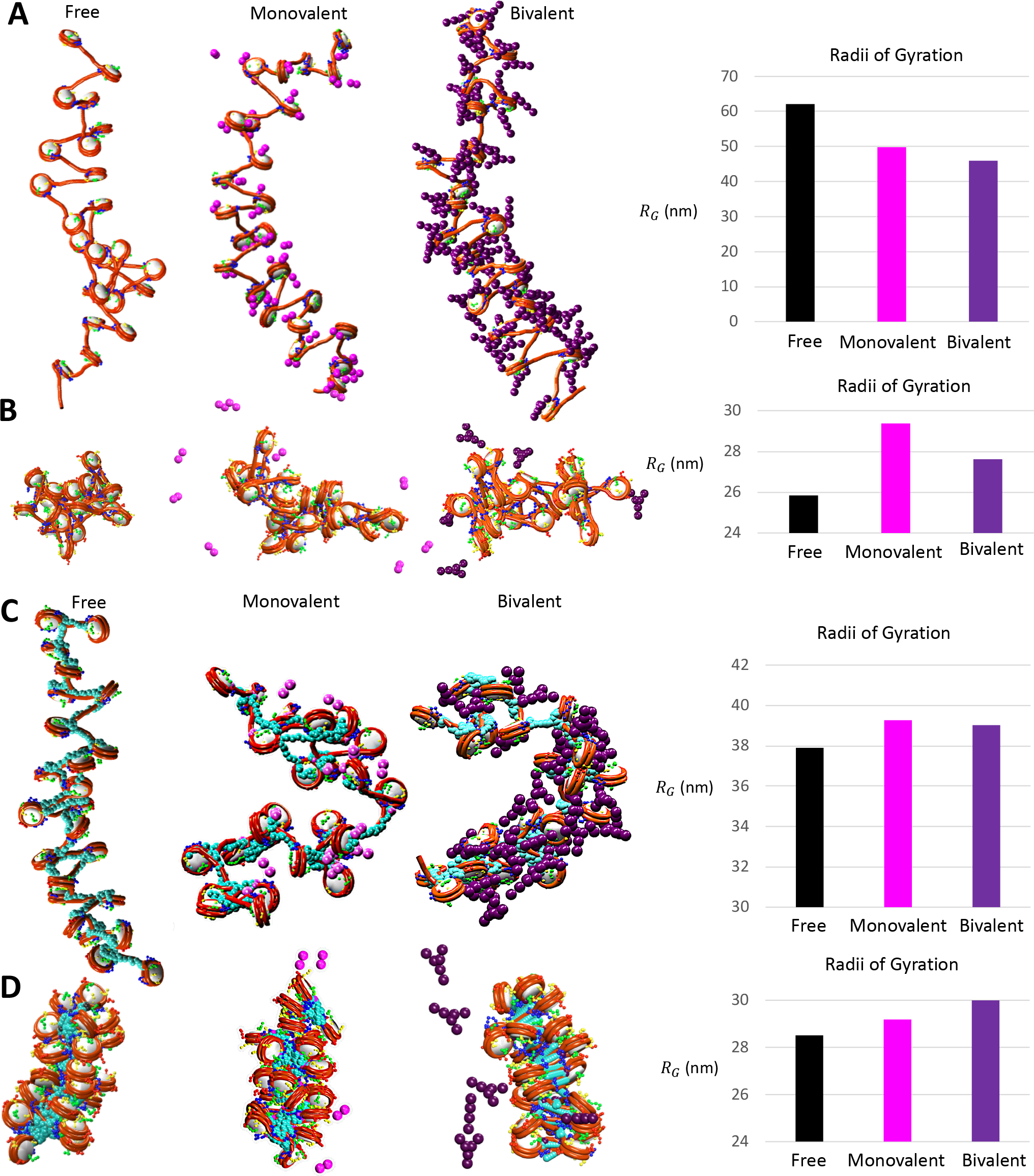
Snapshots of representative fibers from all 12 antibody/chromatin systems. All fibers consist of 24 nucleosomes and start from idealized zigzag configurations (see SI). For each condition (low salt / physiological salt) we show results for free fiber, monovalent Fab complex, and bivalent Fab complex with and without LH. Radii of gyration (*R*_*G*_) of chromatin fibers are shown in nanometers (nm) for free (black), monovalent (magenta), and bivalent (purple) systems. (A) Low salt without LH. (B) Physiological salt without LH. (C) Low salt with LH. (D) Physiological salt with LH.

**Figure 4:**
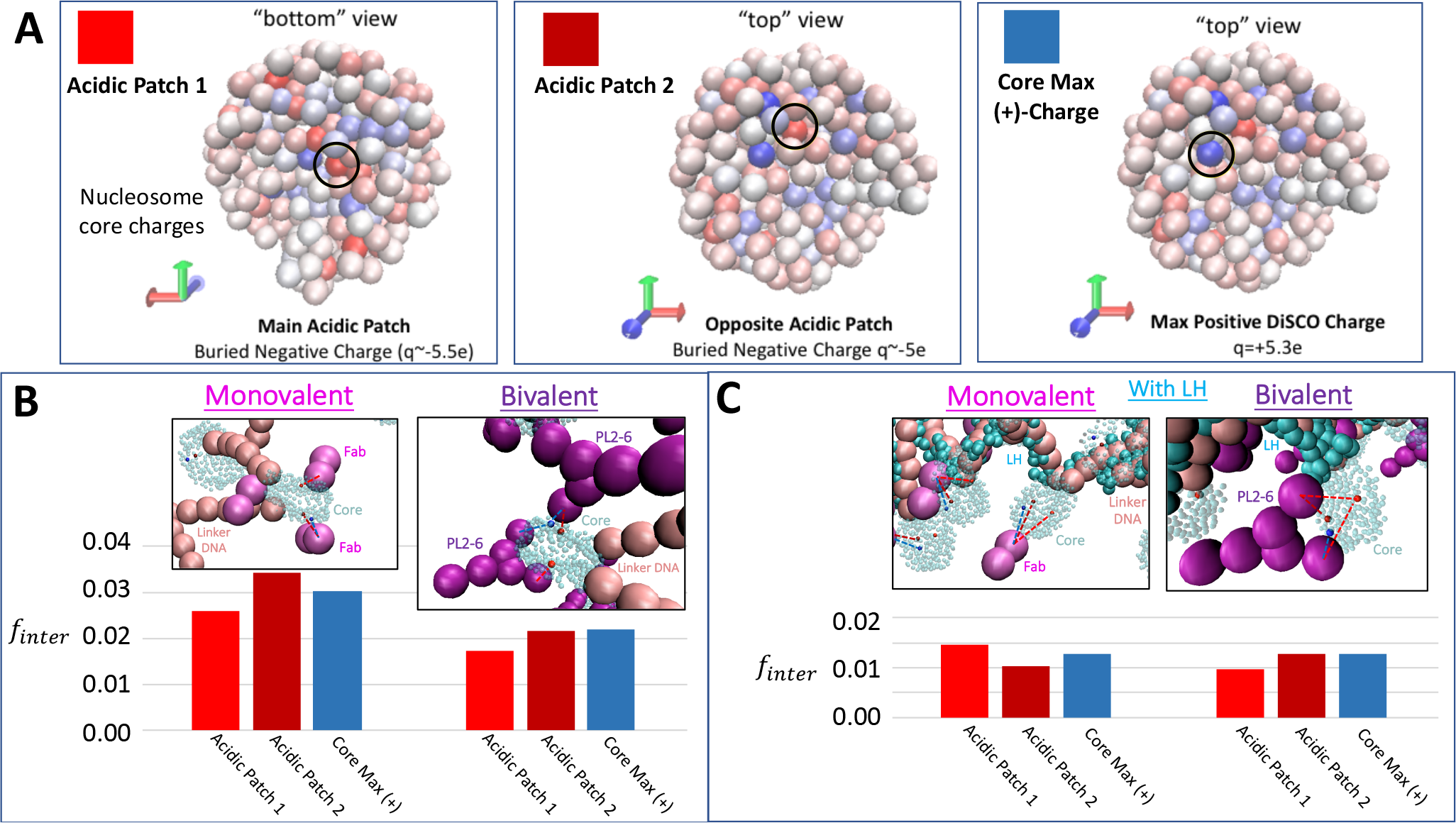
Fab interactions with specific regions of the nucleosome core. Regions include the two negatively-charged acidic patches (arbitrarily labeled as Acidic Patch 1 and 2) and the largest positive charge on the nucleosome surface (core-max-(+)). Interactions (*f*_*inter*_) are evaluated by measuring the number of simulation frames for which pairwise distances between elements are within 5 nm, sampled over the last 1 million frames of the simulation and normalized across all sampled frames. Pairwise distances are measured from the center of the specified charged bead on the nucleosome core and from the center of the charged ‘Fab’ beads in both monovalent and bivalent systems. (A) Low salt without LH. (B) Low salt with LH.

## Simulation Parameters

Idealized zigzag starting structures of 24 nucleosomes and a nucleosome repeat length (NRL, equal to 147 bp plus length of linker DNA) of 200 bp were used for all chromatin fibers, corresponding to a common NRL for human cells (see SI). For simulations containing antibodies, we introduce monovalent and bivalent antibodies at starting configurations distributed, at a distance of 100 nm, evenly along the fiber structure for a total of 72 antibodies (Figures S2 and S3 in SI). To prevent diffusion of the antibody molecules away from the fiber without imposing any artificial forces on the system, we employ periodic boundary conditions (PBC) inside a 400-nm box centered around the chromatin fiber. All simulations were conducted for a minimum of 20 million MC steps.

## Data Analysis

Interactions (*f*_*inter*_) between all chromatin elements and antibodies are evaluated by measuring the number of simulation frames for which pairwise distances between elements are within 5 nm, sampled over the last 1 million frames of the simulation and normalized across all sampled frames. Tail interactions are averaged over both copies of each C-terminal tail for H3, H4, and H2B. *H*2*A*_1_ and *H*2*A*_2_ refer to C-terminal and N-terminal H2A tails, respectively. Distances are measured from the center of the end tail bead, from the geometric center of the entire nucleosome core, from the center of each linker bead, and from the center of the charged ‘Fab’ beads in both monovalent and bivalent systems.

**Figure 5:**
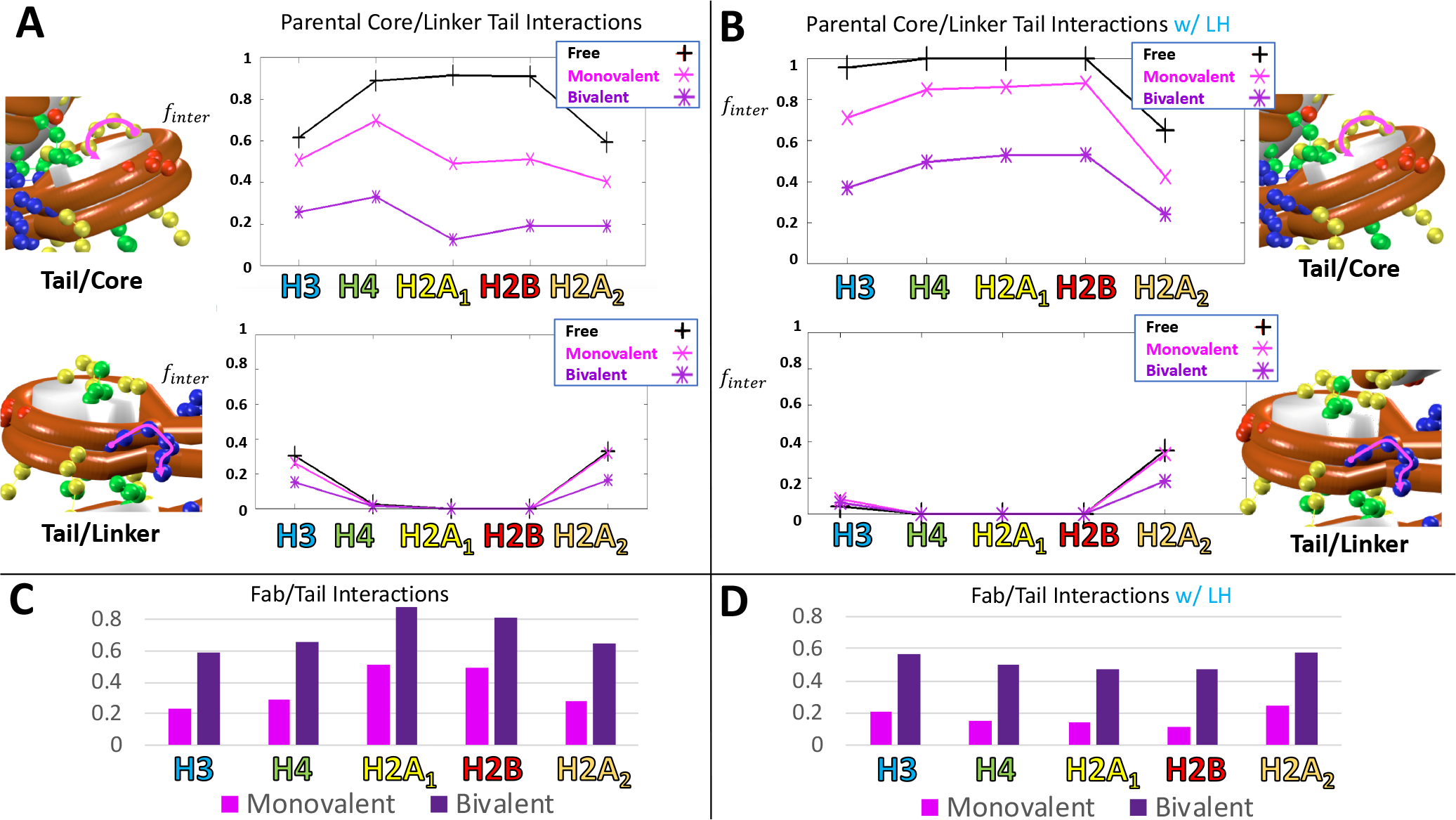
Tail interactions with chromatin and antibodies at low salt. Interactions (*f*_*inter*_) are evaluated by measuring the number of simulation frames for which pairwise distances between elements are within 5 nm, sampled over the last 1 million frames of the simulation and normalized across all sampled frames. Tail interactions are averaged over both copies of each C-terminal tail for H3, H4, and H2B. H2A_1_ and H2A_2_ refer to C-terminal and N-terminal H2A tails, respectively. Distances are measured from the center of the end tail bead, from the geometric center of the entire nucleosome core, from the center of each linker bead, and from the center of the charged ‘Fab’ beads in both monovalent and bivalent systems. (A) Tail interactions with parental cores and parental linkers at low salt without LH. (B) Tail interactions with parental cores and parental linkers at low salt with LH. (C) Tail interactions with antibodies at low salt without LH. (D) Tail interactions with antibodies at low salt with LH.

## RESULTS

We simulate three categories of chromatin systems: **Free** chromatin reference systems, **Monovalent** PL2-6 with chromatin, and **Bivalent** PL2-6 with chromatin. For each of these categories, we simulate at both low and physiological monovalent salt concentrations (*C*_*S*_ = 10 and *C*_*S*_ = 150 mM, respectively), as well as with and without linker histone (LH). Representative structures from each of these 12 systems are shown in Figure 3. For open fiber morphologies under low salt conditions, we observe intense differential binding to chromatin between bivalent vs. monovalent antibodies (Fig. 3A). For condensed fibers at physiological salt, we find that antibody interactions with chromatin fiber are highly dynamic and transient (Fig. 3B). We also show in Fig. 3 the radii of gyration of chromatin fibers for free (black), monovalent (magenta), and bivalent (purple) systems. At low salt (Fig. 3A), both monovalent and bivalent antibodies help to condense chromatin fibers. This behavior resembles synergistic mechanisms of dynamic condensation regulated by LH (**?**). As with LH, positively-charged antibodies compact fibers by neutralizing the negative charge of linker DNA. With bivalent PL2-6’s greater positive charge, we observe further compaction in bivalent compared to monovalent complexes (Fig. 3A,B). Figure 3C shows that, at low salt, this sensitivity to antibody condensation is lost with LH, since condensation due to LH dominates, with or without PL2-6. For chromatin fibers at physiological salt, we see the opposite effect. Since H1-containing chromatin fibers are already more condensed, the addition of antibodies (Fig. 3C,D) increases *R*_*G*_ by disrupting native chromatin contacts. We analyze and discuss these interactions in more detail in the *Physiological Salt* section. In the next section, we focus on “open” chromatin fibers, since they are more statistically reliable in yielding analyses of internal chromatin interactions with monovalent vs. bivalent antibodies to elucidate the causes of differential binding.

**Figure 6:**
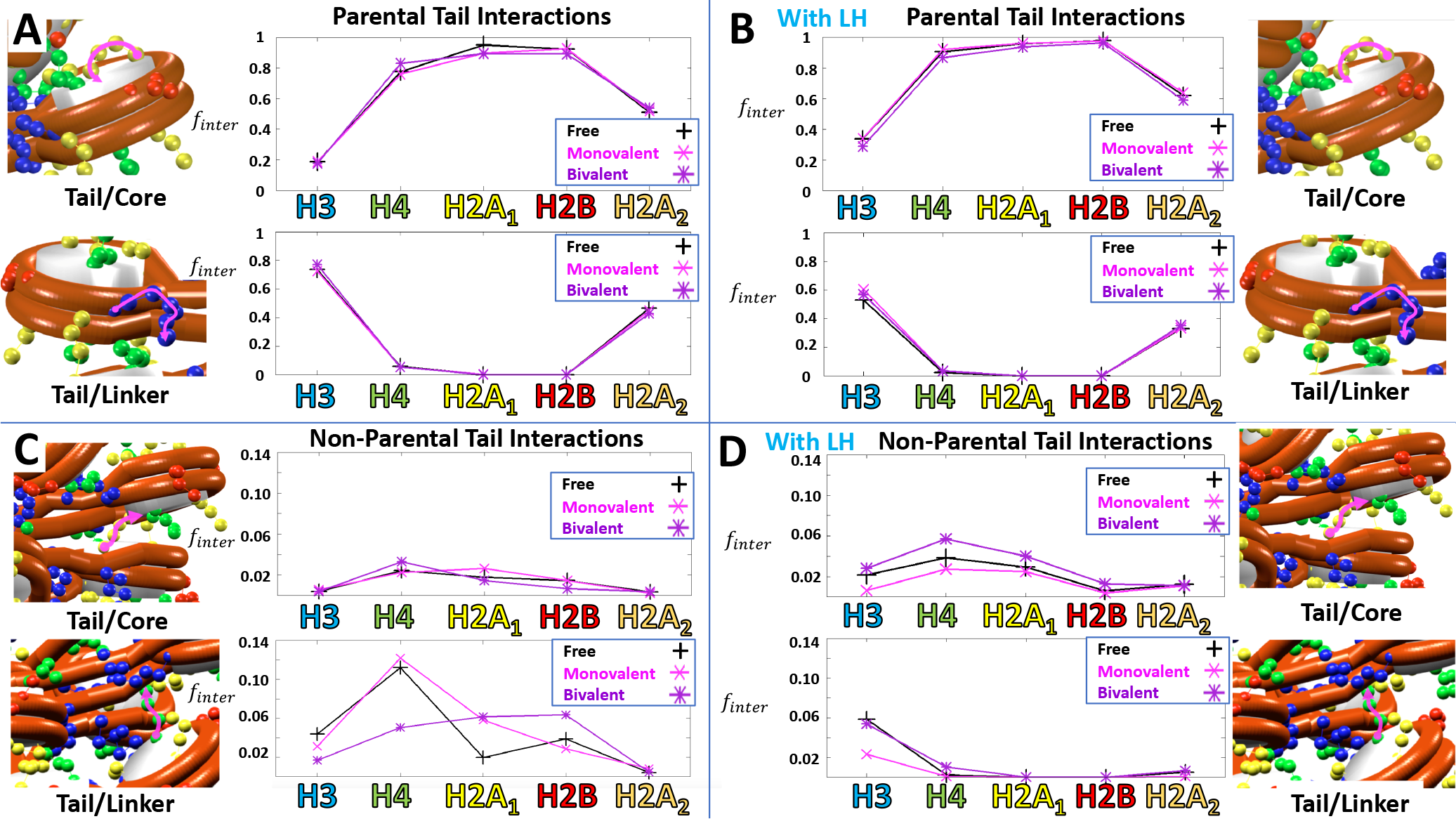
Tail interactions at physiological salt without LH. Interactions (*f*_*inter*_) are evaluated by measuring the number of simulation frames for which pairwise distances between elements are within 5 nm, sampled over the last 1 million frames of the simulation and normalized across all sampled frames. Tail interactions are averaged over both copies of each C-terminal tail for H3, H4, and H2B. H2A_1_ and H2A_2_ refer to C-terminal and N-terminal H2A tails, respectively. Pairwise distances are measured from the center of the end tail bead, from the geometric center of the entire nucleosome core, from the center of each linker bead, and from the center of the charged ‘Fab’ beads in both monovalent and bivalent systems. Tail interactions are shown for both parental and non-parental nucleosome cores and linker DNA for chromatin fibers at physiological salt. (A) Parental tail interactions at physiological salt. (B) Parental tail interactions at physiological salt with LH. (C) Non-parental tail interactions at physiological salt. (D) Non-parental tail interactions at physiological salt with LH.

## Open Zigzag Fibers

### Acidic patch affinity relative to other core nucleosome regions is unaffected by antibody type and LH, but diminishes with bivalent antibody through overall core affinity

We analyze open fibers at low salt to determine whether antibody type and chromatin binding affect the relative acidic patch affinities of PL2-6. Since PL2-6 ‘Fabs’ are expected to bind the acidic patch of mononucleosomes (**?**), it is plausible that differences in chromatin-bound bivalent and monovalent complexes might reflect variations in acidic patch availabilities, which may help account for tighter chromatin binding of bivalent PL2-6. To compare acidic patch affinities between bivalent and monovalent PL2-6 relative to other regions of the nucleosome core, we measure interactions between the Fabs’ binding region (charged spheres) with three selected beads (Fig. 4A, black circles), each representing a separate region on the nucleosome surface: the two acidic patches (on front and back, denoted Acidic Patches 1 and 2 and colored red/dark red), and (as a control) the largest concentration of positive charge on the nucleosome as defined by the charges in our discretized model (blue). The positions of the acidic patches are defined in our model by the center of the most negatively-charged bead on either side of the nucleosome disc. Interactions are measured by counting pairwise distances within a 5 nm cutoff over the final million MC steps of the simulation. From Fig. 4B (without LH), we see no significant differences between the three regions within the same systems of bivalent or monovalent antibodies. However, the overall reduction in core affinity results in a lower acidic patch affinity for *bivalent* PL2-6 compared to *monovalent* Fab fragments without LH. With LH (Fig. 4C), we again note litt difference between these nucleosome core regions. The differences between monovalent and bivalent interactions are also less pronounced in this case. Thus, acidic patch availability does not appear to account for the major observed differential binding of monovalent versus bivalent PL2-6.

**Figure 7:**
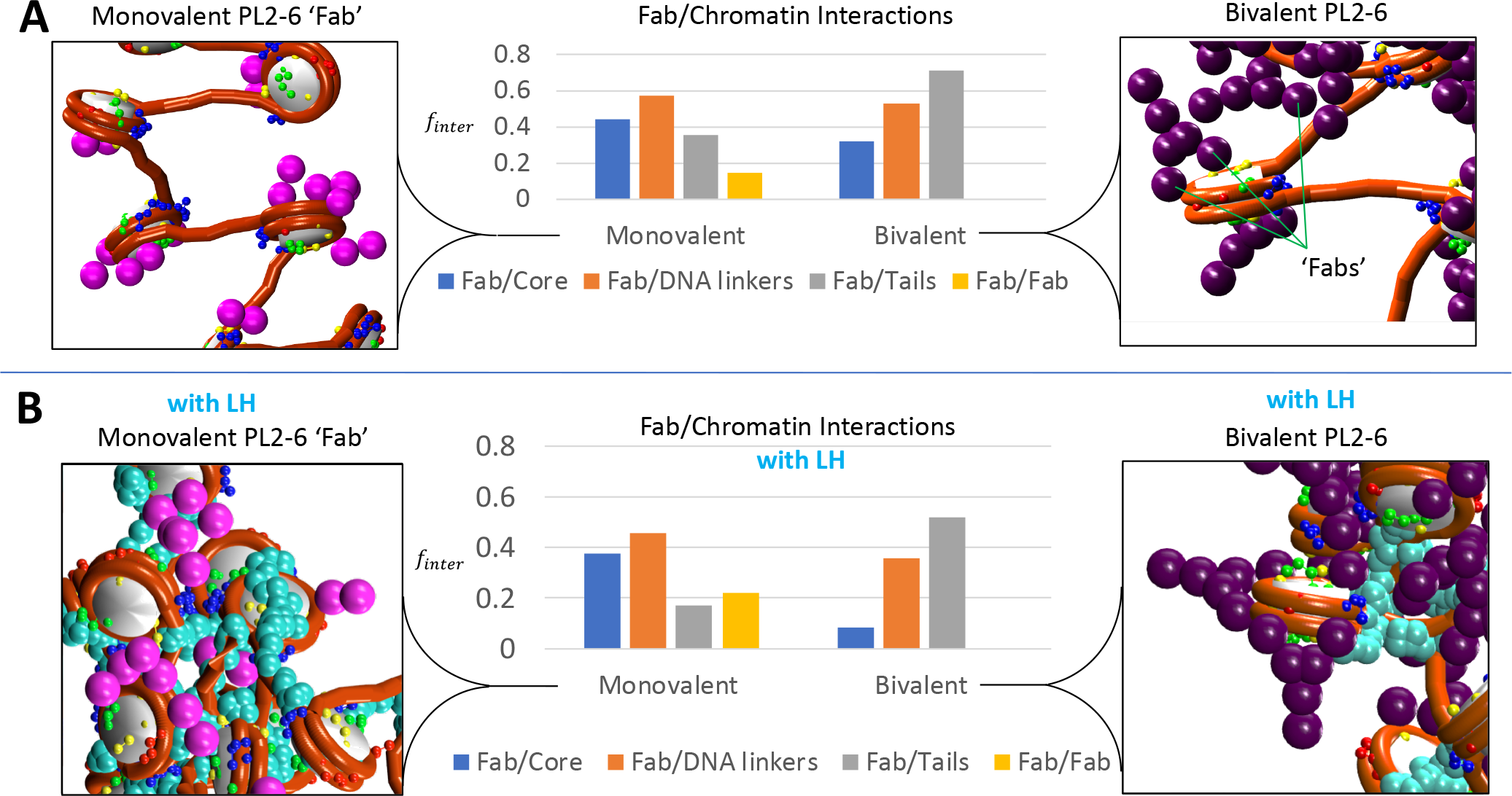
Overall antibody interactions with elements of open chromatin fiber (cores, DNA linkers, tails, and other Fabs) at low salt. Interactions (*f*_*inter*_) are evaluated by measuring the number of simulation frames for which pairwise distances between elements are within 5 nm, sampled over the last 1 million frames of the simulation and normalized across all sampled frames. Interactions for are averaged over all elements in the fiber for each category (core, linkers, tails, Fabs). Pairwise distances are measured from the center of the end tail bead, from the geometric center of the entire nucleosome core, from the center of each linker bead, and from the center of the charged ‘Fab’ beads in both monovalent and bivalent systems. (A) Low salt without LH. (B) Low salt with LH.

### Competition for internal chromatin interactions drives differential antibody binding for bivalent versus monovalent PL2-6 in open chromatin fibers

Next, we look at changes in the internal chromatin structure resulting from the differential binding of bivalent versus monovalent PL2-6 in open chromatin fibers. Figure 5 analyzes interactions for all 6 open chromatin systems between histone tails (H3, H4, H2A_1_, H2B, and H2A_2_) with nonparental cores and nonparental linkers (Fig. parts A,B) and between histone tails and PL2-6 Fab regions (C,D), with (A,C) and without LH (B,D). In Figure 5A, our results for the control system of free chromatin interactions (black) are consistent with previous studies of open, elongated chromatin fibers obtained at low salt (**?**). Such fibers have minimal long-range contacts (non-parental tail/core and tail/linker interactions, not shown) but dominant short-range tail interactions with parental cores (Fig. 5A). Due to their proximity to the entry/exit site along the dyad axis, H3 and N-terminal H2A_2_ tails maintain small interactions with parental linker DNA. With LH (Fig. 5B), linker interactions with H3 tails are reduced through competition with LH for linker DNA contacts, freeing up H3 tails for greater interaction with parental cores in the absence of PL2-6. Figure 5A,C show that these internal chromatin contacts are significantly disrupted by monovalent Fab (magenta) PL2-6 and reduced even more in the presence of bivalent (purple) PL2-6. Similarly, while the tail/linker interactions are not significantly affected by the monovalent (magenta) PL2-6, they are disrupted by bivalent (purple) PL2-6 binding. From Figure 5C,D we see that tails with increased monovalent and bivalent ‘Fab’ interactions correspond to the same tails with decreased internal nonparental core and linker contacts in Figure 5A,B). Thus, differential ‘Fab’ binding of monovalent versus bivalent PL2-6 relies on direct competition with internal fiber/tail contacts. Beyond the electrostatic attraction to negatively-charged linker DNA and regions of the core, the chromatin binding (and thus, differential binding of bivalent vs. monovalent chromatin complexes) is likely driven by entropy gained from increased conformational freedom of histone tails upon binding to freely-moving antibodies.

### Physiological Salt

#### PL2-6 disrupts fiber condensation at physiological salt with monovalent/bivalent differences mediated by LH

Next, we look at the changes in internal chromatin structure for condensed fibers at physiological monovalent salt concentrations. In Figure 6 we analyze tail interactions in the same way as for open fibers (Fig. 5), but also show the longer-range non-parental interactions not present in open fibers. Figure 6 shows interactions for all 6 systems at physiological salt between histone tails (H3, H4, H2A_1_, H2B, and H2A_2_) with parental cores and parental linkers (Fig. parts A,B) and tails with non-parental cores and non-parental linkers (C,D), with (A,C) and without LH (B,D). In the absence of LH (Fig. 6A), tail interactions with parental cores and linkers are similar to those at low salt except that competition between parental linkers and parental cores for H3 and C-terminal H2A_2_ interactions favors the parental linkers, thus decreasing the H3 and C-terminal H2A_2_ contacts with parental cores. In the presence of LH (Fig. 6B), this effect is dampened due to competition between LH and linkers for tail contacts, particularly H3, as it is the closest in proximity to the linker entry/exit site. Without the Fab/tail interactions as seen at low salt, the Fabs provide insufficient competition with internal tail interactions, and the parental core/tail and parental linker/tail interactions remain unchanged with the addition of monovalent or bivalent PL2-6, with or without LH. Chromatin fibers remain satisfied with these internal chromatin contacts, preventing significant binding to either monovalent or bivalent PL2-6. Internucleosomal tail interactions (with non-parental cores and linker DNA), on the other hand, are altered by the presence of monovalent and bivalent PL2-6. Without LH, free fibers are generally more condensed at physiological salt (150 mM) (Fig. 3B). However, this condensation is affected by even minimal interactions with monovalent and bivalent PL2-6. Without LH (Fig. 6C), monovalent PL2-6 increases the stiffness of the fiber (thus increasing *R*_*G*_, Fig. 3B), whereas chromatin with bivalent PL2-6 is more condensed. With LH, free fibers are already stiffer and thus limited in the extent of compaction (Fig. 3D). In contrast, without LH, chromatin with bivalent PL2-6 is less condensed than chromatin with monovalent PL2-6 due to changes in internal chromatin structure. From Fig. 6D, we observe modest but significant histone tail interactions with non-parental cores, enhanced in the presence of bivalent PL2-6, and slightly repressed by monovalent PL2-6. Through competition for linker DNA interactions with LH, tail interactions with non-parental linker DNA are negligible, except for H3 tails, which are closest to the linker DNA. These are left unchanged by bivalent antibodies, whereas the smaller monovalent antibodies are able to transiently penetrate into the stiff fiber structure and disrupt this modest interaction. In short, while monovalent and bivalent PL2-6 interact only transiently with chromatin at physiological salt concentrations, they both alter internal chromatin structure in distinct ways and with LH dependence.

#### Entropic interactions with histone tails a key component in driving differential binding

Finally, to analyze the interactions which cause this differential binding, we plot Fab/chromatin interactions for both antibodies in Fig. 7. Here we see that bivalent antibodies show decreased interactions compared to monovalent system for both core (blue) and linker DNA (orange) but increased tail interactions (gray). Thus, differential binding of bivalent antibodies is driven largely by entropic contributions of increased histone tail/antibody interactions. It is possible that the additional degrees of freedom of the less symmetric rotation axes of bivalent PL2-6 will increase the entropy of histone tails to help promote chromatin binding. The decreased Fab/Fab interactions (yellow) suggest that the increased chromatin binding of bivalent antibodies creates greater disruption of the self-interactions of the system compared to the monovalent complex, as bivalent interactions do not register at the 5 nm cutoff after significant chromatin binding. We capture similar trends with LH (Fig. 7B), but the tail interactions are reduced. Thus, with LH, the competition for tail interactions between PL2-6 and Fab is reduced, making the cores less accessible to Fab binding, particularly in bivalent PL2-6.

## DISCUSSION

By combining mesoscale chromatin and coarse-grained protein models, we demonstrate that changes in internal chromatin structure *in silico* result in differential chromatin binding for bivalent versus monovalent PL2-6 antibodies in open chromatin fibers. We find that differential binding is not affected through acidic patch availability relative to other nucleosome core regions, although acidic patch affinity might affect binding indirectly through changes in overall core affinity. Competition for internal chromatin interactions drives differential antibody binding between bivalent and monovalent PL2-6 in open chromatin fibers. At physiological salt, transient interactions of monovalent and bivalent PL2-6 with chromatin are sufficient to alter the internal chromatin structure and interactions, with LH modulating differences in structure and condensation of chromatin complexes. For open fibers, differences in Fab binding to histone tails play the largest role in driving differential binding, which favors bivalent over monovalent PL2-6 binding. Such modeling and observations should help understand differences in internal chromatin structure which could drive experimentally observed differential binding of bivalent versus monovalent PL2-6.

The structural details that differentiate ‘epichromatin’ from internal chromatin and lead to the observed differential immunostaining patterns remain unclear. In addition, because immunostaining microscopy is conducted on formaldehyde-fixed and detergent permeabilized cells with the antibody incubations performed in physiological PBS buffer (**? ? ?**), molecular motion of the chromatin is somewhat suppressed and space for antibody diffusion may be reduced. Experiments on isolated and unfixed cell nuclei incubated with labeled mono-and bivalent anti-nucleosome antibodies could help further assess the simulation observations. A more comprehensive survey of simulated chromatin structures and conditions with coarse-grained antibody models should provide a better understanding of the observed *in vitro* differential binding and the changes in internal chromatin structure that stabilize such binding. In addition, further refinement of coarse-grained models may provide more insight at physiologically-relevant conditions. Nonetheless, the present work indicates that internal chromatin structure plays a pivotal role in determining differences in antibody binding which can dynamically alter internal chromatin structure. Such feedback mechanisms are expected to be important in other systems involving chromatin binding, such as transcription factors, linker histone, and histone remodeling proteins.

## AUTHOR CONTRIBUTIONS

C.G.M. and T.S. designed the research. C.G.M. performed the simulations and analyzed the data. C.G.M, T.S., D.O., and A.O. wrote the paper.

## Supporting information

Supplemental Text and Figures with Additional Methods

## ACKNOWLEDGMENTS

We thank Robyn Stanfield for providing the PL2-6 ‘Fab’ homology model used for electrostatic calculations. This work was supported by the National Institutes of Health, National Institute of General Medical Sciences awards R01-GM055264 and R35-GM122562, and Phillip-Morris USA and Phillip-Morris International to T.S. DEO and ALO gratefully acknowledge the University of New England, College of Pharmacy for supporting our research, and Bates College and Travis Gould (Department of Physics and Astronomy) for supporting our joint research. Simulations were performed on the New York University high-performance computing (HPC) cluster Prince and the private cluster Schulten.

