## Supplemental Text and Figures with Additional Methods for "Mesoscale Modeling of a Nucleosome-Binding Antibody (PL2-6): Mono- vs. Bivalent Chromatin Complexes"

#### Model Constituents

Histone tails are modeled at a resolution of 5 amino acids per bead. Similarly, LHs are represented as 6 beads for the globular head (GH) and 22 beads for the flexible C-terminal Domain (CTD) as described in Ref. (1). See Ref. (2) for further model details and validation of equilibrium and dynamic properties.

#### Initial Starting Configurations

The initial 3D structure for all chromatin fibers are generated as ideal zigzag conformations with the long fiber axis oriented parallel to the z-axis. The z-rise per nucleosome (distance between successive nucleosomes along the long fiber axis) and fiber width (distance between successive nucleosomes when viewed in the plane perpendicular to the fiber axis) are assigned values proportional to the DNA linker length between nucleosomes such that DNA/DNA linker bead distances are less than the DNA bead radius (3 nm). Any overlaps between cores or linker DNA beads are removed. Nucleosomes are initially oriented perpendicular to the central axis, according to the zigzag structure found to be optimal in Ref. (3) (Fig. S1) due to stabilization by the H3 tail interactions with linker DNA.

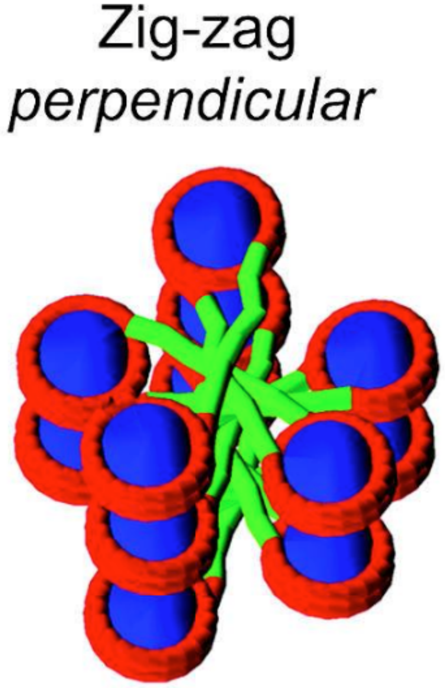


**Figure S1**. Initial perpendicular zigzag chromatin fiber structure (Ref. 3).

Initial coordinates for all monovalent and bivalent antibodies are placed along an approximate cylinder of 100 nm from each nucleosome, with 3 antibodies (each spaced at an additional 20 nm for the second and third antibodies per nucleosome). The first of each antibody triplet per nucleosome is translated 100 nm from the nucleosome center to the antibody center, along the *line of nodes*—defined as the $x'$-axis in the rotated reference frame determined by each nucleosome's initial Euler angle coordinate frame. Each monovalent antibody's long 'Fab' axis is oriented *parallel* with the nucleosome's *line of nodes* (Fig. S2) while each bivalent antibody's long 'Fc' axis is oriented *perpendicular* to the nucleosome's *line of nodes* (Fig. S3).

### Monovalent with Linker Histone (LH)

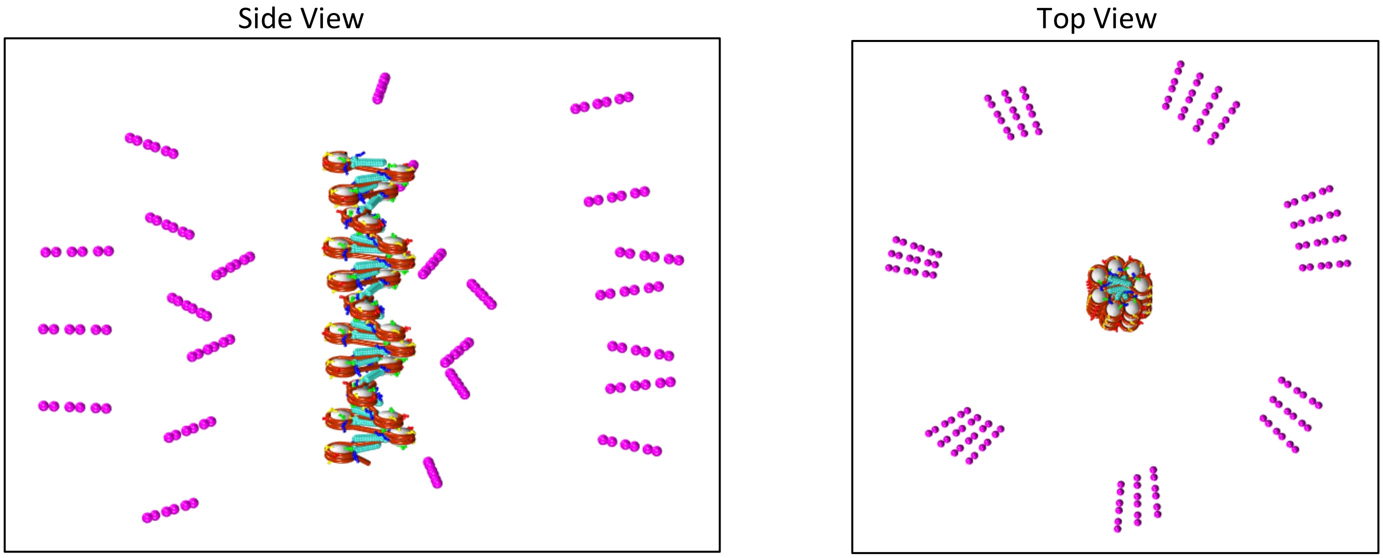


**Figure S2**. Initial positions for zigzag chromatin fibers with monovalent antibodies used in our MC simulations. Representative example with linker histone.

### Bivalent without Linker Histone (LH)

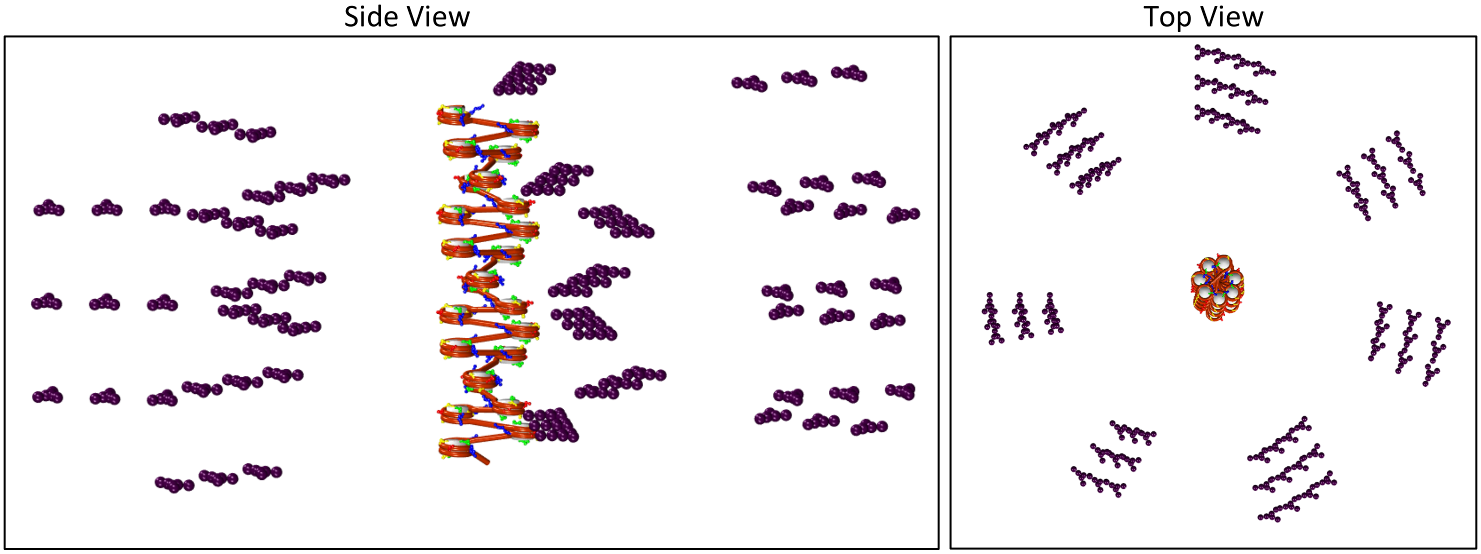


**Figure S3**. Initial positions for zigzag chromatin fibers with bivalent antibodies used in our MC simulations. Representative example without linker histone.

### <u>References</u>:


1. Luque A, Collepardo-Guevara R, Grigoryev S, Schlick T (2014) Dynamic condensation of linker histone C-terminal domain regulates chromatin structure. Nucl. Acids Res. 42:7553–7560.
2. Bascom G, Schlick T (2018) Chromatin fiber folding directed by cooperative histone tail acetylation and linker histone binding. Biophys. J. 114(10):2376–2385.
3. Sun J, Zhang Q, Schlick T (2005) Electrostatic mechanism of nucleosomal array folding revealed by computer simulation. Proc. Natl. Acad. Sci. USA 102:8180—8185.
